# CaATP prolongs strong actomyosin binding and promotes futile myosin stroke

**DOI:** 10.1101/756627

**Authors:** Jinghua Ge, Akhil Gargey, Irina V. Nesmelova, Yuri E. Nesmelov

## Abstract

Calcium plays an essential role in muscle contraction, regulating actomyosin interaction by binding troponin of thin filaments. There are several buffers for calcium in muscle, and those buffers play a crucial role in the formation of the transient calcium wave in sarcomere upon muscle activation. One such calcium buffer in muscle is ATP. ATP is a fuel molecule, and the important role of MgATP in muscle is to bind myosin and supply energy for the power stroke. Myosin is not a specific ATPase, and CaATP also supports myosin ATPase activity. The concentration of CaATP in sarcomeres reaches 1% of all ATP available. Since 294 myosin molecules form a thick filament, naïve estimation gives three heads per filament with CaATP bound, instead of MgATP. We found that CaATP dissociates actomyosin slower than MgATP, thus increasing the time of the strong actomyosin binding. The rate of the basal CaATPase is faster than that of MgATPase, myosin readily produces futile stroke with CaATP. When calcium is upregulated, as in malignant hyperthermia, kinetics of myosin and actomyosin interaction with CaATP suggest that myosin CaATPase activity may contribute to observed muscle rigidity and enhanced muscle thermogenesis.

## Introduction

Calcium plays significant role in regulation of muscle contraction. Calcium binds troponin of thin filaments and initiates its conformational change, leading to opening of myosin binding site in the filament. In muscle, calcium ions are released from the sarcoplasmic reticulum and enter a sarcomere at the Z line (Baylor and Hollingworth 1998). Then, diffusion delivers calcium ions to their binding sites located along thin filaments. Several calcium buffers exist in sarcomere. These buffers mediate calcium transport, producing specific waveform of calcium transient in sarcomere (Baylor and Hollingworth 2012). ATP is one of those buffers. Calcium competes with magnesium to bind ATP, and the resulting transient CaATP concentration in sarcomere reaches 64 - 71 µM (Baylor and Hollingworth 1998; Baylor and Hollingworth 2012). This is about a hundred times less than the concentration of MgATP, which is a fuel molecule for muscle contraction. Considering the number of myosin heads in one thick filament to be equal to 294 heads (Craig and Offer 1976), 1% of available CaATP could lead to several myosin heads in a filament having CaATP bound instead of MgATP. Does this affect muscle contraction? It should, if myosin CaATPase kinetics is different from MgATP kinetics. According to the current hypothesis of force generation in muscle, myosin heads “crawl” on a thin filament, binding to and detaching from actin. During muscle contraction, the duration of the strong actomyosin binding should be short enough to allow multiple myosin heads work together without interference and quickly propel thin filaments. Quick ATP-induced actomyosin dissociation in skeletal muscle was confirmed by measurements of myosin MgATPase (Harris and Warshaw 1993; Uyeda et al. 1990). However, the transient kinetics of actomyosin CaATPase was not studied, leaving several open questions: (a) How fast CaATP binds actomyosin? (b) What is the rate of CaATP-induced actomyosin dissociation? (c) Does CaADP slow down actomyosin dissociation?

The goal of this paper is to examine the kinetics of actomyosin-CaATP interaction, and to predict whether the myosin CaATPase plays a role in muscle contraction. We focused on (a) kinetics of CaATP-induced actomyosin dissociation, and (b) competitive inhibition of actomyosin ATPase by CaADP, because ADP dissociation from actomyosin may determine the time of the strong actomyosin binding during the force-generating step. In order to determine the total duration of the actomyosin cycle, we measured the rate of the steady state actin-activated myosin CaATPase. We found that myosin spends longer time in the strongly bound to actin state with CaATP as a substrate. The longer lasting actin-bound crossbridges should slow muscle contraction. Based on our experimental data we calculated the probability of finding one strongly bound actomyosins in one thick filament. At normal physiological calcium concentration in a sarcomere, the probability of one myosin head be strongly bound to actin is 68% per one thick filament. At pathologic conditions such as malignant hyperthermia (Struk et al. 1998), the calcium release from the sarcoplasmic reticulum in skeletal muscle is substantially upregulated. At increased calcium concentration, the probability to find a strongly bound myosin head increases, which could contribute to the observed muscle rigidity (Ali et al. 2003; Cully et al. 2014).

## Methods

### Reagents

*N*-(1-pyrene)iodoacetamide (pyrene) was from Life Technologies Corporation (Grand Island, NY), phalloidin, ATP, and ADP were from Sigma-Aldrich (Milwaukee, WI). All other chemicals were from ThermoFisher Scientific (Waltham, MA) and VWR (Radnor, PA).

### Proteins preparation

Myosin and actin were prepared from rabbit leg and back muscles (Margossian and Lowey 1982b; Strzelecka-Golaszewska et al. 1980; Waller et al. 1995). Chymotryptic S1 was prepared as described (Waller et al. 1995), and dialyzed into experimental buffer. F-actin was labeled with pyrene iodoacetamide (6:1 label:actin molar ratio), cleaned from the excess label, re-polymerized, concentrated to 200 µM and stabilized with phalloidin at a molar ratio of 1:1, and dialyzed for two days at T=4°C against the experimental buffer. Concentration of unlabeled G-actin was determined spectrophotometrically assuming the extinction coefficient ε_290nm_ = 0.63 (mg/ml)^-1^cm^-1^ (Houk and Ue 1974). Concentration of labeled G-actin and labeling efficiency were determined spectroscopically using the following expressions: [G-actin]=(A_290nm_–(A_344nm_·0.127))/26,600 M^-1^ and [pyrene]=A_344nm_ /22,000 M^-1^ (Takagi et al. 2008). The experimental buffer contained 20 mM MOPS (3-[N-morpholino]propanesulfonic acid) pH 7.3, 50 mM KCl, 0.2 mM EDTA, 0.2 mM EGTA, and 3mM MgCl_2_ or CaCl_2_ total concentration. Since log_10_K_A_ for MgATP and CaATP are 4.29 and 3.91 accordingly (Sigel and Song 1996), where K_A_ is the association constant, 3mM MgCl_2_ or CaCl_2_ chelate all ATP used in our experiments, since used ATP concentration was 1mM or less. We do not expect any measurable effect from the KATP complex, since the association constant for such a complex is three orders of magnitude smaller than constants for CaATP and MgATP (Melchior 1954). All reported concentrations are final concentrations.

### ATPase assays

Steady state basal and actin activated myosin ATPase activities were measured spectrophotometrically at T = 12°C in the buffer contained 20 mM MOPS pH 7.3, 50 mM KCl, and 5mM MgCl_2_ or CaCl_2_, by mixing myosin or actomyosin with 5mM MgATP or CaATP, and monitoring the liberation of inorganic phosphate, as described previously (Lanzetta et al. 1979). Both myosin and F-actin were carefully washed before each experiment to avoid any possibility of contamination by products of hydrolysis, phosphate and ADP. Myosin or actomyosin was mixed with ATP, and aliquots were collected at equal time intervals and analyzed for phosphate in ammonium molybdate - malachite green colorimetric assay (see SI for more details). The rate of basal myosin ATPase V_basal_ was determined as the rate of phosphate production. To determine maximum velocity of the actomyosin ATPase, the rates of myosin ATPase at different actin concentrations were fitted by the hyperbolic equation v = V_basal_ + V_act_[actin]/(K_app_ +[actin]), where v is the rate of the ATPase in the presence of actin, horizontal asymptote, V_max_ = V_basal_ + V_act_, is the rate of myosin ATPase at infinite actin concentration. Normalized values for phosphate concentration at different reaction times (basal ATPase) and reaction rates at different actin concentrations (actin activated ATPase) obtained from experiments using several independent preparations of actin and myosin were averaged and mean values and standard deviations were plotted and fitted with the line (basal ATPase) or with the hyperbola (actin activated ATPase) using Origin 8 (OriginLab Corp., Northampton MA). Determined ATPase rates (the value and the standard error) are presented in the Table 1. We did not account for F-actin ATPase activity since this is a slow process (Martonosi et al. 1960), significantly inhibited by phalloidin (Dancker et al. 1975).

**Table 1.**
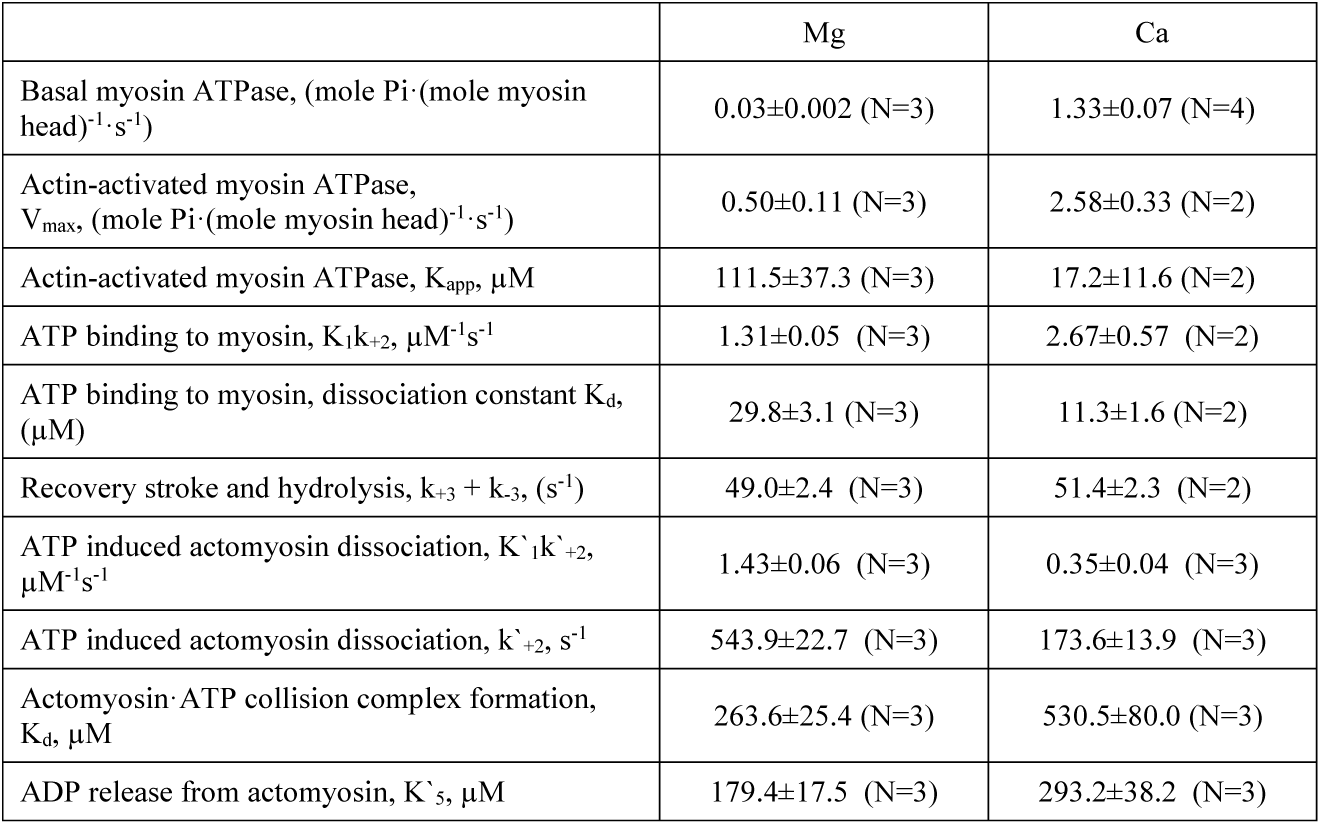
Myosin and actomyosin kinetic rate constants for magnesium and calcium nucleotide complexes. The data are averages of N independent protein preparations

### Acquisition of fluorescence transients

Transient intrinsic tryptophan fluorescence of myosin and fluorescence of pyrene actin were measured with Bio-Logic SFM-300 stopped flow transient fluorimeter (Bio-Logic Science Instruments SAS, Claix, France), equipped with FC-15 cuvette. The mixing unit dead time is 2.6 ms (Figure S4). All experiments were performed at T=12° C to reliably detect transient kinetics of actomyosin dissociation by ATP, at higher temperature the rate becomes too fast for reliable detection (Figure S3). Usually, three syringes and two mixers were used in an experiment. The continuous flow and the smallest intermixer delay line (40 μL) were usually used. Myosin intrinsic fluorescence was excited by mercury-xenon lamp at 296 nm and detected using a 320 nm cutoff filter. The pyrene fluorescence was excited at 365 nm and detected using a 420 nm cutoff filter. Multiple transients were usually acquired and averaged to improve signal to noise ratio. 8000 points were acquired in each experiment.

### Analysis of fluorescence transients

Transients obtained in each experiment were fitted globally by the single exponential function S(t) = S_o_+A·exp(-k_obs_·(t-t_0_)), or the double exponential function S(t) = S_o_+A_1_·exp(-k_obs1_·(t-t_0_)) + A_2_·exp(-k_obs2_·(t-t_0_)). S(t) is the observed signal at time t, A_i_ is the signal amplitude, t_0_ is the time when flow stops (Figure S4), and k_obsi_ is the observed rate constant. To determine the bimolecular rate (K_1_k_+2_ and K’_1_k’_+2_, the asterisk indicates reaction rate constants describing actomyosin kinetics), the dependence of the observed rates on the nucleotide or protein concentration was fitted by a straight line at small concentrations of the nucleotide. Competitive inhibition of ATP-induced actomyosin dissociation by ADP was measured with pyrene labeled actin complexed with myosin and incubated with ADP at various concentrations, including [ADP] = 0 µM. This solution was rapidly mixed with ATP, and the transient fluorescence of pyrene-actin dissociated from myosin was measured. The observed reaction rates were fitted globally to the equation k_obs_ = V_max_·[ATP]/(K_d_·(1 + [ADP]/K’_5_)+[ATP]) (Deacon et al. 2012; Rosenfeld et al. 2000; Segel 1976), with several common parameters: rate constant V_max_ (k’_+2_), the equilibrium constant of actomyosin·ATP collision complex dissociation K_d_ (1/K’_1_), and K’_5_, the equilibrium constant of ADP dissociation from actomyosin upon chasing with ATP (Figure 1). Rate constants obtained from experiments with proteins of usually three independent preparations were averaged and mean values and standard deviations were used to fit with equations listed above. Determined constants are the mean value and the standard error. All data fits were performed with Origin 8 (OriginLab Corp, Northampton MA). Statistical significance of results were tested with ANOVA integrated in Origin 8 software. A significance level of P < 0.05 was used for all analyses.

**Figure 1.**
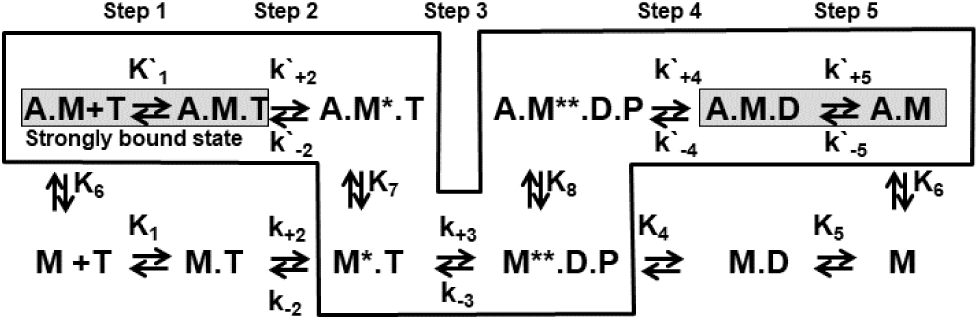
Acto-myosin ATPase cycle reaction scheme. A = actin, M = myosin (M* and M** - myosin states with increased intrinsic fluorescence, fluorescence in M** state is higher), T = ATP, D = ADP, P = phosphate. Asterisk indicates reaction rate constants describing actomyosin kinetics. Highlighted = strongly bound actomyosin state. Boxed = generally accepted pathway of actomyosin interaction.

## Results

### The rates of basal and actin-activated myosin CaATPase activity are faster than for MgATP

It is well known that actin activates myosin MgATPase. Actin activation is defined as the increased rate of myosin ATPase activity in the presence of actin. Our data show about 20-fold increase of actin activation of myosin MgATPase activity, in excellent agreement with previous studies (Lymn and Taylor 1971; Rosenfeld and Taylor 1987). The measured rates of basal MgATPase and actin activated MgATPase are 0.03 ± 0.002 mole Pi·(mole myosin head)^-1^·s^-1^ (Figure 2), and 0.47 ± 0.11 mole Pi·(mole myosin head)^-1^·s^-1^ (Figure 3), respectively. The measured rate of basal CaATPase and actin activated CaATPase are 1.33±0.07 mole Pi·(mole myosin head)^-1^·s^-1^ (Figure 2), and 1.25±0.33 mole Pi·(mole myosin head)^-1^·s^-1^ (Figure 3). Note that the reaction rate of actin activated CaATPase observed in experiment (Figure 3) is the sum of rates of basal and actin-activated ATPases, as explained in the Methods. Measured rates of basal MgATPase and CaATPase agree well with the published results (Bagshaw and Trentham 1974; Banerjee and Morkin 1978; Shriver and Sykes 1981; Sleep et al. 1981), the rate of basal CaATPase is 50 times higher than the rate of MgATPase. The value of actin activation (V_max_/V_basal_) of myosin CaATPase is just two-fold, smaller than the value for MgATPase (20-fold), in agreement with the previous report (Polosukhina et al. 2000). Measured rate of actin activated MgATPase is smaller than the rate, reported in literature for low temperatures. This is because the literature data were obtained with buffers of low ionic strength, which increases the rate of actin activated myosin ATPase. We measured actin activated myosin MgATPase at low ionic strength ([KCl]=0) and found the rate constant 1.54 ± 0.07 mole Pi·(mole myosin head)^-1^·s^-1^, in good agreement with published results for actin activated myosin MgATPase at low temperature and in low ionic strength buffer (Travers and Hillaire 1979; White et al. 1993).

**Figure 2.**
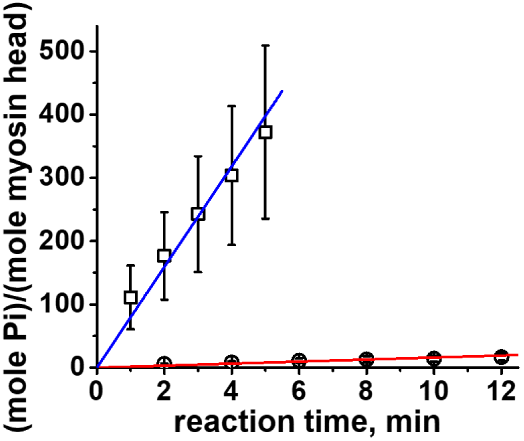
Basal myosin ATPase activity, [myosin] = 3.3 μM, [ATP] = 5 mM. Circles, MgATP, squares, CaATP. Linear fit. The data points are averages of N = 3 (Mg) and N=4 (Ca) independent protein preparations. Uncertainties are SD here and throughout the text. v = 0.03±0.002 mole Pi·(mole myosin head)^-1^·s^-1^ (Mg), and 1.33±0.07 mole Pi·(mole myosin head)^-1^·s^-1^ (Ca).

**Figure 3.**
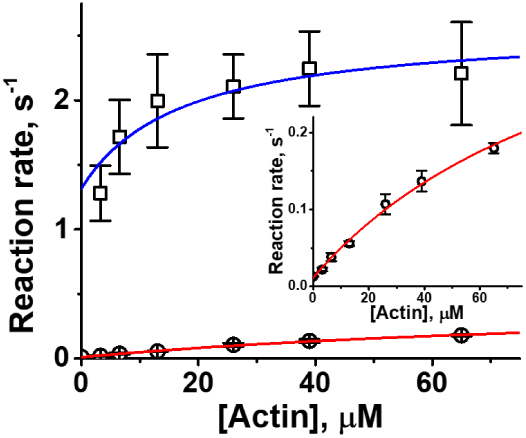
Steady state actin activated myosin ATPase activity, [myosin] = 0.8 μM, [ATP] = 5 mM. Circles, MgATP, squares, CaATP. Line, fit with a hyperbola, V_basal_ + V_max_[actin]/(K_app_ +[actin]). Insert, MgATPase. The data points are averages of N=3 (Mg) and N=2 (Ca) independent protein preparations. V_max_ (basal ATPase activity is subtracted) is 0.50±0.11 mole Pi·(mole myosin head)^-1^·s^-1^, (Mg), and 2.58±0.33 mole Pi·(mole myosin head)^-1^·s^-1^ (Ca).

### The effect of metal cation on myosin-ATP interaction

The rate of ATP-induced myosin conformational change was measured using 0.5 µM myosin S1 (here and throughout the text the concentration in the final mixture is given), rapidly mixed with various concentrations of MeATP, where Me = Mg or Ca. Myosin intrinsic fluorescence changes upon mixing with ATP. Observed transients were fitted with one (Mg) or two (Ca) exponentials (Figure 4). Both metal cations sustain myosin ATPase activity, showing the increase of M** population (myosin high fluorescence state) upon mixing myosin S1 with ATP. First order reaction rate constants of ATP binding to myosin, K_1_k_+2_, are 1.31±0.05 μM^-1^s^-1^ for MgATP (in excellent agreement with previous reports (Bagshaw 1975; Bagshaw and Trentham 1973; Lymn and Taylor 1971)) and 2.67±0.57 μM^-1^s^-1^ for CaATP. Maximum rate of myosin conformational change V_max_ is the same within experimental error for both cations, 49.0±2.4 s^-1^ for MgATP and 51.4±2.3 s^-1^ for CaATP (Figure 5). This rate is slower than previously determined due to lower temperature in our experiments. Figure 4 shows that in the presence of MgATP, myosin S1 reaches the long-lasting steady state M**. The upper limit of the M** lifetime is 33 s, or 1/V_max_ of basal myosin MgATPase. Our data also show that in the presence of CaATP, myosin reaches the M** state. However, the M** state depopulates with the rate 0.99±0.08 s^-1^, and the equilibrium is established at the lower level of myosin intrinsic fluorescence. Similar effect was observed before for rabbit skeletal myosin MnATPase (Bagshaw 1975) and D. discoideum myosin (Tkachev et al. 2013). This behavior indicates that the equilibrium between the M** and M* states is shifted towards the M* state in the presence of CaATP. The rate of depopulation of the M** state does not depend on CaATP concentration. Therefore, the lifetime of the M** state in myosin CaATPase is a constant value, which is shorter than the lifetime of the M** state in myosin MgATPase. The rate of the M** state depopulation in myosin CaATPase corresponds well to 1/V_max_ of the basal myosin CaATPase. We interpret observed depopulation of M** state in the presence of calcium as faster Pi release, followed by ADP release. Slow ADP release should result in deeper drop of myosin intrinsic fluorescence in Figure 4, not observed in our experiments. We exclude weak CaATP binding to myosin in interpretation of observed transient kinetics, since CaATP binds tighter to myosin than MgATP (K_app Mg_ > K_app Ca_, Table 1), and observed depopulation of M** state does not depend on ATP concentration. Myosin lifetime in the M** state in the presence of calcium is long enough for myosin head to bind the thin filament, since CaATP in muscle is at maximum at high calcium, when the highest number of binding sites on thin filament is available.

**Figure 4.**
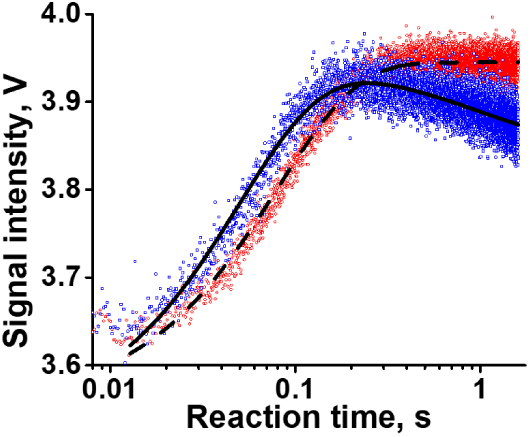
Myosin intrinsic fluorescence transients upon rapid mixing of myosin and ATP. Myosin 0.5 μM, MeATP 10 μM. Me=Mg, red dots, experiment, and dashed line, single exponential fit. Me=Ca, blue dots, experiment, and solid line, double exponential fit. The second exponent in the case of CaATP does not depend on [CaATP], the rate is 0.99±0.08 s^-1^.

**Figure 5.**
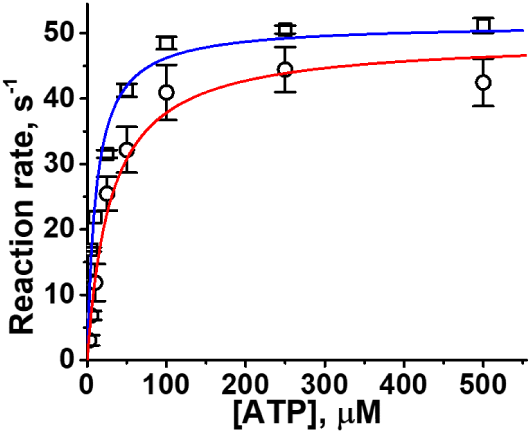
Kinetics of ATP binding and recovery stroke upon rapid mixing of myosin and ATP; observed rate constants are fitted with a hyperbola, V_max_ = 49.0±2.4 s^-1^ for MgATP (circles, N=3) and V_max_ = 51.4±2.3 s^-1^ for CaATP (squares, N=2).

### The rate of CaATP-induced actomyosin dissociation is three times slower than that for MgATP

It is known that ADP does not limit the ATP-induced dissociation of rabbit skeletal actomyosin, and there is a fast equilibrium between actomyosin and actomyosin·ADP complex (Geeves 1989; Siemankowski et al. 1985; Woodward et al. 1991). In this case, the maximum rate of the ATP-induced actomyosin dissociation does not depend on the competitive inhibition by ADP (Segel 1976). In our experiments, 0.5 μM of pyrene-labeled actomyosin was premixed and equilibrated with MeADP and then rapidly mixed with MeATP (Me = Mg or Ca). All observed transients of pyrene fluorescence were well-fitted with a single-exponential function, confirming the rapid-exchange equilibrium of free ADP and ADP bound to actomyosin (De La Cruz and Ostap 2009). The following concentrations of ADP and ATP were used in the experiment: ADP, 0 µM, 50 µM, 100 µM, 250 µM, 500 µM, 750 µM, 1 mM, and 1.25 mM; ATP, 25 µM, 50 µM, 100 µM, 250 µM, 500 µM, 1 mM, 2 mM, and 2.5 mM. To obtain the reaction rates of actomyosin dissociation at different ADP and ATP concentrations, obtained transients were fitted globally, keeping the parameters k’_+2_, K_d_, and K’_5_ the same for all transients, using the model of competitive inhibition (Segel 1976) (Figure 7). The global fit decreases the error in the determination of kinetic parameters of interest. We found that the maximum rate of the ATP-induced actomyosin dissociation (k’_+2_) is almost three times smaller for CaATP than for MgATP (Table 1). The first order reaction rate constant K’_1_k’_+2_ is 1.43±0.06 µM^-1^s^-1^ for MgATP, in excellent agreement with literature data (Criddle et al. 1985; Marston and Taylor 1980). For CaATP the rate constant K’_1_k’_+2_ is four times smaller, 0.35±0.04 µM^-1^s^-1^ (Figure 6). The equilibrium constant of the formation of the collision complex actomyosin·CaATP (K’_1_ = 1/K_d_) is two times higher than for MgATP (Table 1). The equilibrium constant of ADP dissociation from actomyosin (K’_5_) is 60% higher for CaADP (Table 1). The data for magnesium are in a good agreement with published results (Geeves 1989; Nyitrai et al. 2006; Siemankowski et al. 1985). Prepared actomyosin·MeADP complexes have had the same level of pyrene fluorescence, that confirms strong actomyosin binding (De La Cruz and Ostap 2009) in the presence of magnesium and calcium.

**Figure 6.**
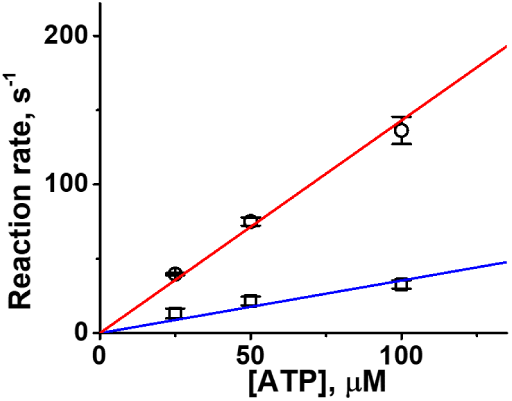
ATP-induced actomyosin dissociation. Observed reaction rates at low [ATP] are fitted by a straight line, the second order reaction rate constant is determined from the slope of the line. K’_1_k’_+2_ = 1.43±0.06 µM^-1^s^-1^, MgATP, and 0.35±0.04 µM^-1^s^-1^, CaATP. Circles, MgATP, N=3, squares, CaATP, N=3.

**Figure 7.**
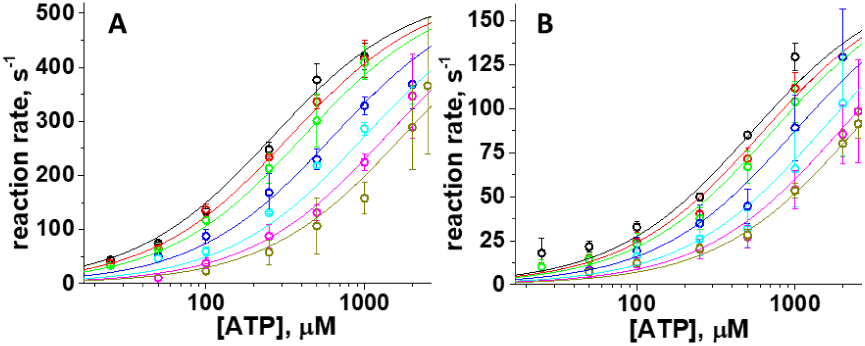
ATP-induced actomyosin dissociation, competitive inhibition with ADP. Global fit of data for multiple ATP and ADP concentrations, listed in the text. Observed reaction rates are fitted with a hyperbola, V_max_ = 543.9±22.7 s^-1^, MgATP (A), and 173.6±13.9 s^-1^, CaATP (B). N=3. Note that y axes are different in A and B panels.

## Discussion

Muscle contraction is a well-orchestrated process that involves multiple proteins and small molecules, in which timing is highly important for the whole operation. According to the current model of force generation in muscle, the short time of the strong actomyosin binding is critical for the synergistic work of multiple myosin heads on one thin filament. CaATP increases the time of strong binding because the rate of CaATP-induced actomyosin dissociation is slower than the rate for MgATP. This increased duration of strong binding should affect the force generation in muscle.

A similar effect is observed in frictional loading assays, when various actin binding proteins or NEM myosin bind to actin, providing frictional load that opposes the propelling force, produced by myosin, and slows down the actin filament mobility (Greenberg and Moore 2010).

When the supply of ATP is unlimited and myosin binding sites of the thin filament are available, the probability that one myosin head is strongly bound is the ratio of the time of the strongly bound state to the time of the actomyosin cycle. MeADP release from actomyosin is fast for rabbit skeletal myosin for magnesium (Geeves 1989; Siemankowski et al. 1985) and calcium, since we observe only single-exponential transients in all range of ADP concentrations. If the rate of ADP release is slower or compared to the rate of ATP induced actomyosin dissociation, we should have two-exponential transients, especially at ADP concentrations, close to the equilibrium constant of ADP dissociation K’_5_ (De La Cruz and Ostap 2009). Therefore, assuming k’_+5_ > k’_+2_, where k’_+5_ is the rate of ADP release from actomyosin, the rate of ATP-induced actomyosin dissociation determines the time of the strong binding, and t_on_ ∼ 1/k’_+2_, instead of t_on_ = 1/k’_+2_ + 1/k’_+5_. This is the upper limit of the rate of ADP dissociation from actomyosin. The time of the actomyosin cycle is determined from the rate of the steady state actin-activated myosin ATPase, t_total_ = 1/V_max_. Therefore, using data in Table 1, for one myosin head the probability to be strongly bound to actin is P_1Mg_ = (t_on_/t_total_)_Mg_ = (V_max_/k’_+2_)_Mg_ = 9.2·10^−4^ ± 2.1·10^−4^ in the presence of MgATP, and P_1Ca_ = (t_on_/t_total_)_Ca_ = (V_max_/k’_+2_)_Ca_ = 14.9·10^−3^ ± 2.2·10^−3^ in the presence of CaATP. The concentration of MgATP is much higher than the concentration of CaATP in muscle, how does this affect the total probability to find a strongly bound myosin? According to Baylor et al. (Baylor and Hollingworth 1998; Baylor and Hollingworth 2012), the concentration of MgATP in sarcomere is 7.27 mM, and the peak concentration of CaATP is 64 - 71 µM. The concentration of myosin in muscle can be estimated as 113 µM, assuming that each thick filament in a sarcomere occupies the volume of 4.32·10^6^ nm^3^. Indeed, the hexagonal arrangement of thick filaments with 27.5 nm distance between thick and thin filament (Kawai et al. 1993) gives the 1964.8 nm^2^ cross-section of muscle per thick filament. The cross section multiplied by the sarcomere length of 2.2 µm gives the volume of the muscle, occupied by one thick filament. Since 294 myosin molecules form one thick filament, this number of molecules in the volume 4.32·10^6^ nm^3^ corresponds to myosin concentration of 113 µM. Knowing equilibrium constant for the formation of collision actomyosin·ATP complex K’_1_ (K’_1_ = 1/K_d_, Table 1), one can calculate the concentration of such collision complexes, actomyosin·CaATP and actomyosin·MgATP, at the peak concentration of CaATP in muscle (the equation to calculate the concentration of collision complex is derived in the SI). The fraction of actomyosin·CaATP complex is about 10% (it depends on the number of myosin heads strongly bound, from 10.8% for just one strongly bound head in the thick filament to 9.4% when all heads in the thick filament are in the strongly bound state). One can think of this fraction as the probability for actomyosin to form actomyosin·MeATP collision complex, these probabilities are P_2Mg_=0.9 and P_2Ca_=0.1 for Mg and Ca, respectively. The product of the probability to form actomyosin·MeATP complex P_2Me_ (Me = Mg or Ca) and the probability to find myosin in the strongly bound state at unlimited ATP P_1Me_ gives the probability for one myosin head to be in the strongly bound state at the physiological ATP concentration. Since both MgATP and CaATP present in muscle, the total probability to find one myosin head, strongly bound to actin, is the sum of two probabilities, P = P_1Mg_ P_2Mg_ + P_1Ca_ · P_2Ca_. For one myosin head such probability is P = 2.3·10^−3^ ± 0.3·10^−3^ and, since there are 294 myosin heads in one thick filament, the probability that one myosin head in thick filament is strongly bound to actin in thin filament is P_strong_ = P · 294 = 0.68 ± 0.09, or 68%. At elevated calcium concentration (3-fold) the probability that CaATP binds actomyosin increases (P_2Mg_ = 0.78 and P_2Ca_ = 0.22), therefore the probability to find strongly bound myosin head in one thick filament increases to P_strong_ = 1.18 ± 0.15, or 118%. If k’_+5_ is close to k’_+2_ for both ions, magnesium and calcium, which is the lower limit of the rate of ADP dissociation from actomyosin, t_on_ = 1/k’_+2_ + 1/k’_+5_ ∼ 2/k’_+2_. Then, the probability to find a strongly bound crossbridge in a thick filament at physiological calcium is 34%, and at 3-fold elevated calcium the probability increases to 59%. Nearly 50% of myosin heads can be in the super-relaxed state in skeletal muscle (Hooijman et al. 2011), that decreases the concentration of active myosin heads in sarcomere two times, and therefore, the probability to find a strongly bound crossbridge.

While at normal conditions CaATP exists in muscle in small quantity, the disturbance of the skeletal muscle calcium homeostasis, or malignant hyperthermia (MH) (MacLennan and Phillips 1992), results in the elevated calcium concentration in the myoplasm (Ali et al. 2003; Rosenberg et al. 2007). In MH patients, the peak of intracellular calcium release rate is three times greater than in healthy individuals (Struk et al. 1998). MH is the result of the increased permeability of the sarcoplasmic reticulum (SR) membrane and decreased buffering capacity in the SR itself (Manno et al. 2013). Typical signs of MH include muscle rigidity and elevated body temperature. It is hypothesized that the elevated body temperature is the result of thermogenesis due to the increased MgATP consumption by myosin, caused by troponin upregulation, and by SERCA pumping the excess of calcium into SR (MacLennan and Phillips 1992). Muscle rigidity is explained by depletion of ATP due to its increased consumption (Rosenberg et al. 2007). Our observations of myosin and actomyosin kinetics in the presence of calcium show that myosin readily produces futile reverse recovery stroke with CaATP (Figure 4), with myosin basal CaATPase 50 times faster than MgATPase. We propose that fast and futile myosin CaATPase might be one of the reasons of excessive ATP consumption at elevated levels of calcium in muscle, leading to thermogenesis. Increased probability to have strongly bound myosin head due to increased concentration of CaATP may contribute to observed muscle rigidity, even when ATP in muscle is not depleted.

The rate of ATP binding to actomyosin is close to the rate of the diffusion-controlled reaction (Ge et al. 2016). Our PFG NMR studies show similar coefficients of CaATP and MgATP diffusion in buffered solution (Figure S1), indicating that the diffusion of CaATP does not affect its interaction with actomyosin. Previous studies showed similar rates of myosin·CaADP and myosin·MgADP binding to actin (Tkachev et al. 2013). The pyrene-labeled actomyosin has the same intensity of pyrene fluorescence in the presence of CaADP and MgADP, indicating similar strong binding of actomyosin in the presence of either calcium or magnesium cations. Therefore, the effect of slow actomyosin dissociation by CaATP cannot be minimized by unavailability of CaATP due to not adequate diffusion, or the absence of the strongly bound actomyosin in the presence of CaADP.

The probability of strong binding of myosin head to thin filament is the duty ratio, and this value, determined in our study for myosin with MgATP as a substrate, is lower than the generally accepted value for skeletal myosin (0.04-0.05 (Harris and Warshaw 1993; Uyeda et al. 1990)). We explain this discrepancy as the inherent difference of methods used to determine the duty ratio. There are several methods to determine the duty ratio: (a) as the ratio of the number of actively stroking myosin heads to the total number of myosin heads, capable of interaction with actin filament (Fusi et al. 2017; Harada et al. 1990; Harris and Warshaw 1993; Uyeda et al. 1990; Webb et al. 2013), (b) as the ratio of the times that myosin spends in the strongly bound state (t_s_) to the total time of the actomyosin cycle (t_c_) (Heissler et al. 2013; Sommese et al. 2013), and (c) as the mole fraction of myosin in the strongly bound state in the presence of ATP (De La Cruz et al. 2000; Guhathakurta et al. 2017; Henn and De La Cruz 2005). The last two methods were used to determine the duty ratio of non-muscle myosin II (Heissler et al. 2013), bovine cardiac myosin (Guhathakurta et al. 2017), and myosin V (De La Cruz et al. 2000) and VII (Henn and De La Cruz 2005). The value of the duty ratio for skeletal myosin (0.04 – 0.05) was determined using the former method from the analysis of the sliding velocity of actin filament in the in vitro motility assay (Harris and Warshaw 1993; Uyeda et al. 1990), and from the analysis of muscle fiber compliance at unloaded shortening and data on thick filament spacing in X-ray diffraction experiments (Fusi et al. 2017). The ADP release from actomyosin can be slower in the in vitro motility assay as compared to myosin kinetics in solution due to the internal strain within myosin molecule and potential effect of neighboring motors (Yengo et al. 2012). Mechanical strains affect duration of the strongly bound state (Finer et al. 1994; Harris and Warshaw 1993). Therefore, the duty ratio determined from the in vitro motility data cannot be compared to the duty ratio determined in solution studies, especially when dilute solutions are used, as in our work. Other factors that can affect the duty ratio include temperature, at which the data are collected, and the ionic strength of solution. The rate of ADP release from actomyosin is less temperature dependent than the rate of actin-activated myosin ATPase (Siemankowski et al. 1985). Therefore, myosin duty ratio should increase with temperature. The rate of actin-activated myosin ATPase depends on the solution ionic strength, and the ATPase activity is frequently measured at low ionic strength to obtain maximal rate of actomyosin interaction (Margossian and Lowey 1982a; Polosukhina et al. 2000; Rosenfeld and Taylor 1984; Rosenfeld and Taylor 1987; White et al. 1993). The duty ratio is higher, when t_s_ is determined at high ionic strength and t_c_ at low ionic strength (Sommese et al. 2013). In this work, we conducted our experiments at low temperature and kept the ionic strength of buffered solutions in all experiments the same. Despite of smaller value of myosin duty ratio, determined in our studies, the important conclusion is that in the presence of calcium, myosin duty ratio is an order of magnitude higher than for magnesium at the same experimental conditions, and even small amounts of calcium modulate average myosin duty ratio.

## Conclusion

ATP is a buffer for calcium, and small amount of CaATP always exists in muscle. Myosin is not a specific ATPase and hydrolyzes CaATP with the same rate as MgATP. Myosin CaATPase is much faster than MgATPase, potentially leading to the futile actomyosin cycle, which may contribute to muscle thermogenesis. Slower rate of CaATP-induced actomyosin dissociation may lead to muscle rigidity due to the extended time of the strongly bound state of actomyosin. For the physiologic concentration of calcium in muscle, the estimated probability to find one strongly bound myosin head per thick filament is 68%. When calcium homeostasis of skeletal muscle is disturbed, the probability to find strongly bound myosin head increases, which may disturb muscle contraction such as in malignant hyperthermia.

### Compliance with ethical standards

Myosin and actin were produced from rabbit skeletal tissue. All experimental protocols were approved by the Institutional Animal Care and Use Committee of UNC Charlotte and all experiments were performed in accordance with relevant guidelines and regulations.

## Acknowledgements

This work was supported by National Institutes of Health grant HL132315 and by funds provided by the University of North Carolina at Charlotte.

## Conflict of interest

The authors declare that they have no conflicts of interest with the contents of this article.

## Reference List

Ali SZ, Taguchi A, Rosenberg H (2003) Malignant hyperthermia Best Pract Res Clin Anaesthesiol 17:519–533

Bagshaw CR (1975) Kinetic Mechanism of Manganous Ion-Dependent Adenosine-Triphosphatase of Myosin Subfragment 1 Febs Lett 58:197–201 doi:Doi 10.1016/0014-5793(75)80258-4

Bagshaw CR, Trentham DR (1973) The reversibility of adenosine triphosphate cleavage by myosin Biochem J 133:323–328

Bagshaw CR, Trentham DR (1974) The characterization of myosin-product complexes and of product-release steps during the magnesium ion-dependent adenosine triphosphatase reaction Biochem J 141:331–349

Banerjee S, Morkin E (1978) Thermodynamic studies on the binding of adenosine diphosphate and calcium to beef cardiac myosin Biochim Biophys Acta 536:10–17

Baylor SM, Hollingworth S (1998) Model of sarcomeric Ca2+ movements, including ATP Ca2+ binding and diffusion, during activation of frog skeletal muscle J Gen Physiol 112:297–316 doi:DOI 10.1085/jgp.112.3.297

Baylor SM, Hollingworth S (2012) Intracellular calcium movements during excitation-contraction coupling in mammalian slow-twitch and fast-twitch muscle fibers J Gen Physiol 139:261–272 doi:10.1085/jgp.201210773

Craig R, Offer G (1976) Axial arrangement of crossbridges in thick filaments of vertebrate skeletal muscle J Mol Biol 102:325–332

Criddle AH, Geeves MA, Jeffries T (1985) The use of actin labelled with N-(1-pyrenyl)iodoacetamide to study the interaction of actin with myosin subfragments and troponin/tropomyosin Biochem J 232:343–349

Cully TR, Edwards JN, Launikonis BS (2014) Activation and propagation of Ca2+ release from inside the sarcoplasmic reticulum network of mammalian skeletal muscle J Physiol 592:3727–3746

Dancker P, Low I, Hasselbach W, Wieland T (1975) Interaction of Actin with Phalloidin - Polymerization and Stabilization of F-Actin Biochim Biophys Acta 400:407–404 doi:Doi 10.1016/0005-2795(75)90196-8

De La Cruz EM, Ostap EM (2009) Kinetic and equilibrium analysis of the myosin ATPase Methods Enzymol 455:157–192

De La Cruz EM, Wells AL, Sweeney HL, Ostap EM (2000) Actin and light chain isoform dependence of myosin V kinetics Biochemistry-Us 39:14196–14202

Deacon JC, Bloemink MJ, Rezavandi H, Geeves MA, Leinwand LA (2012) Erratum to: Identification of functional differences between recombinant human alpha and beta cardiac myosin motors Cell Mol Life Sci 69:4239–4255 doi:10.1007/s00018-012-1111-5

Finer JT, Simmons RM, Spudich JA (1994) Single myosin molecule mechanics: piconewton forces and nanometre steps Nature 368:113–119

Fusi L et al. (2017) Minimum number of myosin motors accounting for shortening velocity under zero load in skeletal muscle J Physiol 595:1127–1142

Ge J, Bouriyaphone SD, Serebrennikova TA, Astashkin AV, Nesmelov YE (2016) Macromolecular Crowding Modulates Actomyosin Kinetics Biophys J 111:178–184

Geeves MA (1989) Dynamic interaction between actin and myosin subfragment 1 in the presence of ADP Biochemistry-Us 28:5864–5871

Greenberg MJ, Moore JR (2010) The molecular basis of frictional loads in the in vitro motility assay with applications to the study of the loaded mechanochemistry of molecular motors Cytoskeleton (Hoboken) 67:273–285 doi:10.1002/cm.20441

Guhathakurta P, Prochniewicz E, Roopnarine O, Rohde JA, Thomas DD (2017) A Cardiomyopathy Mutation in the Myosin Essential Light Chain Alters Actomyosin Structure Biophys J 113:91–100

Harada Y, Sakurada K, Aoki T, Thomas DD, Yanagida T (1990) Mechanochemical coupling in actomyosin energy transduction studied by in vitro movement assay J Mol Biol 216:49–68

Harris DE, Warshaw DM (1993) Smooth and skeletal muscle myosin both exhibit low duty cycles at zero load in vitro The Journal of biological chemistry 268:14764–14768

Heissler SM, Liu X, Korn ED, Sellers JR (2013) Kinetic characterization of the ATPase and actin-activated ATPase activities of Acanthamoeba castellanii myosin-2 The Journal of biological chemistry 288:26709–26720

Henn A, De La Cruz EM (2005) Vertebrate myosin VIIb is a high duty ratio motor adapted for generating and maintaining tension The Journal of biological chemistry 280:39665–39676

Hooijman P, Stewart MA, Cooke R (2011) A New State of Cardiac Myosin with Very Slow ATP Turnover: A Potential Cardioprotective Mechanism in the Heart Biophys J 100:1969–1976 doi:10.1016/j.bpj.2011.02.061

Houk TW, Jr., Ue K (1974) The measurement of actin concentration in solution: a comparison of methods Anal Biochem 62:66–74 doi:0003-2697(74)90367-4 [pii]

Kawai M, Wray JS, Zhao Y (1993) The Effect of Lattice Spacing Change on Cross-Bridge Kinetics in Chemically Skinned Rabbit Psoas Muscle-Fibers .1. Proportionality between the Lattice Spacing and the Fiber Width Biophys J 64:187–196 doi:Doi 10.1016/S0006-3495(93)81356-0

Lanzetta PA, Alvarez LJ, Reinach PS, Candia OA (1979) An improved assay for nanomole amounts of inorganic phosphate Anal Biochem 100:95–97

Lymn RW, Taylor EW (1971) Mechanism of Adenosine Triphosphate Hydrolysis by Actomyosin Biochemistry-Us 10:4617-& doi:DOI 10.1021/bi00801a004

MacLennan DH, Phillips MS (1992) Malignant hyperthermia Science 256:789–794

Manno C et al. (2013) Altered Ca2+ concentration, permeability and buffering in the myofibre Ca2+ store of a mouse model of malignant hyperthermia J Physiol 591:4439–4457 doi:10.1113/jphysiol.2013.259572

Margossian SS, Lowey S (1982a) Preparation of myosin and its subfragments from rabbit skeletal muscle Methods Enzymol 85 Pt B:55–71

Margossian SS, Lowey S (1982b) Preparation of myosin and its subfragments from rabbit skeletal muscle Methods Enzymol 85 Pt B:55–71

Marston SB, Taylor EW (1980) Comparison of the myosin and actomyosin ATPase mechanisms of the four types of vertebrate muscles J Mol Biol 139:573–600

Martonosi A, Gouvea MA, Gergely J (1960) Studies on Actin .1. Interaction of C-14-Labeled Adenine Nucleotides with Actin J Biol Chem 235:1700–1703

Melchior NC (1954) Sodium and potassium complexes of adenosinetriphosphate: equilibrium studies The Journal of biological chemistry 208:615–627

Nyitrai M, Rossi R, Adamek N, Pellegrino MA, Bottinelli R, Geeves MA (2006) What limits the velocity of fast-skeletal muscle contraction in mammals? J Mol Biol 355:432–442 doi:10.1016/j.jmb.2005.10.063

Polosukhina K, Eden D, Chinn M, Highsmith S (2000) CaATP as a substrate to investigate the myosin lever arm hypothesis of force generation Biophys J 78:1474–1481

Rosenberg H, Davis M, James D, Pollock N, Stowell K (2007) Malignant hyperthermia Orphanet J Rare Dis 2:21 doi:10.1186/1750-1172-2-21

Rosenfeld SS, Taylor EW (1984) The ATPase mechanism of skeletal and smooth muscle acto-subfragment 1 The Journal of biological chemistry 259:11908–11919

Rosenfeld SS, Taylor EW (1987) The Mechanism of Regulation of Actomyosin Subfragment-1 Atpase J Biol Chem 262:9984–9993

Rosenfeld SS et al. (2000) Kinetic and spectroscopic evidence for three actomyosin:ADP states in smooth muscle J Biol Chem 275:25418–25426 doi:10.1074/jbc.M002685200

Segel IH (1976) Biochemical Calculations. Wiley,

Shriver JW, Sykes BD (1981) Phosphorus-31 nuclear magnetic resonance evidence for two conformations of myosin subfragment-1.nucleotide complexes Biochemistry-Us 20:2004–2012

Siemankowski RF, Wiseman MO, White HD (1985) ADP dissociation from actomyosin subfragment 1 is sufficiently slow to limit the unloaded shortening velocity in vertebrate muscle Proc Natl Acad Sci U S A 82:658–662

Sigel H, Song B (1996) Solution studies of nucleotide-metal ion complexes - isomeric equilibria. In: Astrid Sigel HS (ed) Metal Ions in Biological Systems, vol 32. CRC Press, p 135

Sleep JA, Trybus KM, Johnson KA, Taylor EW (1981) Kinetic-Studies of Normal and Modified Heavy-Meromyosin and Subfragment.1. J Muscle Res Cell M 2:373–399 doi:Doi 10.1007/Bf00711966

Sommese RF et al. (2013) Molecular consequences of the R453C hypertrophic cardiomyopathy mutation on human beta-cardiac myosin motor function Proc Natl Acad Sci U S A 110:12607–12612

Struk A, Lehmann-Horn F, Melzer W (1998) Voltage-dependent calcium release in human malignant hyperthermia muscle fibers Biophys J 75:2402–2410

Strzelecka-Golaszewska H, Prochniewicz E, Nowak E, Zmorzynski S, Drabikowski W (1980) Chicken-gizzard actin: polymerization and stability Eur J Biochem 104:41–52

Takagi Y, Yang Y, Fujiwara I, Jacobs D, Cheney RE, Sellers JR, Kovacs M (2008) Human myosin Vc is a low duty ratio, nonprocessive molecular motor J Biol Chem 283:8527–8537 doi:10.1074/jbc.M709150200

Tkachev YV, Ge J, Negrashov IV, Nesmelov YE (2013) Metal cation controls myosin and actomyosin kinetics Protein Sci 22:1766–1774

Travers F, Hillaire D (1979) Cryoenzymological studies on myosin subfragment 1. Solvent, temperature and pH effects on the overall reaction Eur J Biochem 98:293–299

Uyeda TQ, Kron SJ, Spudich JA (1990) Myosin step size. Estimation from slow sliding movement of actin over low densities of heavy meromyosin J Mol Biol 214:699–710

Waller GS, Ouyang G, Swafford J, Vibert P, Lowey S (1995) A minimal motor domain from chicken skeletal muscle myosin The Journal of biological chemistry 270:15348–15352

Webb M, Jackson DR, Jr., Stewart TJ, Dugan SP, Carter MS, Cremo CR, Baker JE (2013) The myosin duty ratio tunes the calcium sensitivity and cooperative activation of the thin filament Biochemistry-Us 52:6437–6444

White HD, Belknap B, Jiang W (1993) Kinetics of binding and hydrolysis of a series of nucleoside triphosphates by actomyosin-S1. Relationship between solution rate constants and properties of muscle fibers The Journal of biological chemistry 268:10039–10045

Woodward SKA, Eccleston JF, Geeves MA (1991) Kinetics of the Interaction of 2’(3’)-O-(N-Methylanthraniloyl)-Atp with Myosin Subfragment-1 and Actomyosin Subfragment-1 - Characterization of 2 Acto.S1.Adp Complexes Biochemistry-Us 30:422–430 doi:DOI 10.1021/bi00216a017

Yengo CM, Takagi Y, Sellers JR (2012) Temperature dependent measurements reveal similarities between muscle and non-muscle myosin motility J Muscle Res Cell M 33:385–394

