## Supplementary Information for "CaATP prolongs strong actomyosin binding and promotes futile myosin stroke"

**Steady state ATPase assays**

Concentration of ATP in both basal and actin activated myosin ATPase was 5 mM, myosin concentration was 3.3 μM in the basal and 0.84 μM in the actin activated myosin ATPase. Concentrations between 3.25 μM to 65 μM actin were used in the actin activated ATPase assays. At time zero myosin or actomyosin was mixed with ATP, and aliquots were collected at equal time intervals. The solution was incubated at T = 12°C. After collection, the aliquots were immediately mixed with the malachite green – ammonium molybdate solution (0.06% w/v malachite green carbinol hydrochloride, 15 mM ammonium molybdate tetrahydrate in 1N HCl, with 4% v/v of 1% Sterox detergent, added right before use), the mixture was quenched with 34% w/v sodium citrate in 1N HCl after 30 seconds of color development. Then the aliquots were incubated 20 minutes at room temperature before measurement of optical density at 620 nm. To interpret results of optical density measurement at 620 nm in terms of phosphate concentration, a calibration curve was measured using the same colorimetric assay with premixed phosphate solutions of known concentration. Obtained data of optical density were fitted with the straight lines, to determine the rate of color development in the course of ATPase, then the rates were recalculated in terms of concentration of the phosphate produced in the reaction, using the calibration curve. Normalization on the amount of myosin in the assay gives the rate of the phosphate production per myosin head. Attention was paid to be sure that obtained data fit with the straight lines, to confirm that the presence of ADP, produced in the reaction, does not perturb the reaction. 5 mM ATP is clearly a saturating concentration, and the velocity of the reaction is not affected by ADP contamination.

**PFG NMR diffusion measurements**

PFG NMR diffusion measurements were performed at David H. Murdock Research Institute (Kannapolis, NC) on a Bruker Avance-III 950 MHz spectrometer using a pulsed field gradient spin-echo experiment (ledbpgp2s pulse sequence). The 1D spectra were recorded at T=12ºC with number of scans varied from 16 (1000 μM sample) to 64 (250 μM sample) for each gradient step. The PFG strength (100% = 55.7 G $cm^{-1}$) was increased linearly between 2% and 95% over 12 steps. For unrestricted diffusion of a molecule in an isotropic liquid, the diffusion attenuation of spin echo amplitude, $A(g^2)$, is described by the following equation:

$$A(g^2) = A(0) \exp(- \gamma^2 \delta^2 g^2 t_d D), \qquad \text{Eq.S1}$$

where $A(0)$ is the spin echo amplitude at $g = 0$, $\gamma$ is the gyromagnetic ratio for protons, and $t_d = \Delta - \delta/3$ is the diffusion time. The diffusion coefficient D was determined from the linear slope of the attenuation presented using a semi-logarithmic scale according to Eq. S1. For these experiments, the duration of gradient pulses δ of 4 ms and a fixed diffusion time of 50 ms was used.

**Diffusion coefficients of MgATP and CaATP are similar**

For bimolecular reactions, even in a macroscopically homogeneous solution, the relative diffusion of the reactants is the key step that can influence the reaction rate. Therefore, we first measured the diffusion coefficients of CaATP and MgATP at conditions used in our experiments, T=12°C in buffered aqueous solution. The diffusion coefficient was measured at several different concentrations by using the pulsed field gradient (PFG) NMR technique (**Figure S 1**). We found that the diffusion coefficients of CaATP and MgATP are similar within the experimental error and equal $2.6 \pm 0.1 \cdot 10^{-10}$ $m^2/s$ and $2.7 \pm 0.1 \cdot 10^{-10}$ $m^2/s$ for MgATP and CaATP, respectively. Therefore, any difference in CaATP and MgATP binding to actomyosin is related to the selectivity of the myosin active site.

**Concentration of the collision complex actomyosin·ATP**

In the ATP-induced actomyosin dissociation experiment, hyperbolic fit of actomyosin dissociation rates at different ATP concentration produces constant $K_{app} = 1/K`_1$, where $K`_1$ is the equilibrium association constant formation of collision actomyosin·ATP complex.

We used determined constants $K`_1$ for actomyosin and MgATP and CaATP and calculated the concentration of collision actomyosin·ATP complex, using following derivation.

$[AM]_0 = [AM] + [AM \cdot T]$, where $[AM]_0$ is the initial concentration of actomyosin, $[AM]$ and $[AM \cdot T]$ are concentrations of actomyosin and actomyosin·ATP complex in mixture,
$[T]_0 = [T] + [AM \cdot T]$, where $[T]_0$ is the initial concentration of ATP, $[T]$ and $[AM \cdot T]$ are concentrations of ATP and actomyosin·ATP complex in mixture,
$K_{app} = ([AM] \cdot [T])/[AM \cdot T]$, where $K_{app}$ is the dissociation constant of the collision actomyosin·ATP complex, $1/K`_1$.
Then, $K_{app} = \left(([AM]_0 - [AM \cdot T]) \cdot ([T]_0 - [AM \cdot T])\right)/[AM \cdot T]$, this equation is rearranged into the quadratic equation $[AM \cdot T]^2 - [AM \cdot T] \cdot \left([AM]_0 + [T]_0 + K_{app}\right) + [AM]_0 \cdot [T]_0 = 0$, with a meaningful solution

$$[AM \cdot T] = \frac{\left(([AM]_0+[T]_0+K_{app}) - \sqrt{([AM]_0+[T]_0+K_{app})^2 - 4 \cdot [AM]_0 \cdot [T]_0}\right)}{2}.$$

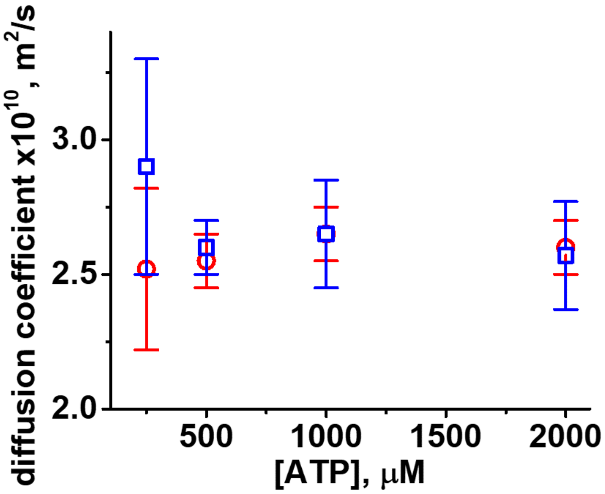


**Figure S 1** Concentration dependence of ATP diffusion coefficients in the presence of Ca and Mg in buffered aqueous solution. T=12°C. Circles, MgATP, squares, CaATP. Within the experimental error, no difference in the diffusion coefficient of CaATP and MgATP is observed.

**Concentration of CaATP at the peak of calcium transient in sarcomere**

Following Baylor and Hollingworth [1], transient concentration of CaATP in sarcomere can be estimated from following kinetic schemes and corresponding equations, taking into account transients of calcium buffers, such as troponin, parvalbumin, and ATP.

Troponin: $Ca + Tr \underset{k_{-1}}{\overset{k_{+1}}{\rightleftarrows}} Ca \cdot Tr$

Parvalbumin: $Ca + Par \underset{k_{-2}}{\overset{k_{+2}}{\rightleftarrows}} Ca \cdot Par \quad and \quad Mg + Par \underset{k_{-3}}{\overset{k_{+3}}{\rightleftarrows}} Mg \cdot Par$

ATP: $Ca + ATP \underset{k_{-4}}{\overset{k_{+4}}{\rightleftarrows}} Ca \cdot ATP \quad and \quad Mg + ATP \underset{k_{-5}}{\overset{k_{+5}}{\rightleftarrows}} Mg \cdot ATP$

Corresponding equations were solved numerically with Wolfram Mathematica 10.3,

$$\frac{d[Tr]}{dt} = -k_{+1}[Ca] \cdot [Tr] + k_{-1}[Ca \cdot Tr]$$

$$\frac{d[Ca \cdot Tr]}{dt} = k_{+1}[Ca] \cdot [Tr] - k_{-1}[Ca \cdot Tr]$$

$$\frac{d[Par]}{dt} = -k_{+2}[Ca] \cdot [Par] + k_{-2}[Ca \cdot Par] - k_{+3}[Mg] \cdot [Par] + k_{-3}[Mg \cdot Par]$$

$$\frac{d[Ca \cdot Par]}{dt} = k_{+2}[Ca] \cdot [Par] - k_{-2}[Ca \cdot Par]$$

$$\frac{d[Mg \cdot Par]}{dt} = k_{+3}[Mg] \cdot [Par] - k_{-3}[Mg \cdot Par]$$

$$\frac{d[ATP]}{dt} = -k_{+4}[Ca] \cdot [ATP] + k_{-4}[Ca \cdot ATP] - k_{+5}[Mg] \cdot [ATP] + k_{-5}[Mg \cdot ATP]$$

$$\frac{d[Ca \cdot ATP]}{dt} = k_{+4}[Ca] \cdot [ATP] - k_{-4}[Ca \cdot ATP]$$

$$\frac{d[Mg \cdot ATP]}{dt} = k_{+5}[Mg] \cdot [ATP] - k_{-5}[Mg \cdot ATP]$$

$$\frac{d[Ca]}{dt} = -k_{+4}[Ca] \cdot [ATP] + k_{-4}[Ca \cdot ATP] - k_{+2}[Ca] \cdot [Par] + k_{-2}[Ca \cdot Par] - k_{+1}[Ca] \cdot [Tr] + k_{-1}[Ca \cdot Tr]$$

We used kinetic constants and initial concentrations from Baylor and Hollingworth [1]:

$k_{+1}$ = 88.5 $\mu M^{-1}s^{-1}$

$k_{-1}$ = 115 $s^{-1}$

$k_{+2}$ = 41.7 $\mu M^{-1}s^{-1}$

$k_{-2}$ = 0.5 $s^{-1}$

$k_{+3} = 0.033\ \mu M^{-1}s^{-1}$

$k_{-3} = 3\ s^{-1}$

$k_{+4} = 13.64\ \mu M^{-1}s^{-1}$

$k_{-4} = 30000\ s^{-1}$

$k_{+5} = 1.5\ \mu M^{-1}s^{-1}$

$k_{-5} = 150\ s^{-1}$

$[ATP]_0$ = 728 µM, (9.1% of 8 mM, total [ATP])

$[Mg{\cdot}ATP]_0$ = 7.27 mM (90.9% of 8 mM, total [ATP])

$[Ca{\cdot}ATP]_0 = 0$

$[Ca]_0$ = 351 µM in the "normal" muscle [1], this parameter was a variable

$[Mg]_0 = [Mg]$ = 1 mM, constant concentration

$[Par]_0$ = 73.5 µM (4.9% of 1.5 mM)

$[Ca{\cdot}Par]_0$ = 615 µM (41% of 1.5 mM)

$[Mg{\cdot}Par]_0$ = 811.5 µM (54.1% of 1.5 mM)

$[Tr]_0$ = 222.9 (92.9% of 240 µM)

$[Ca{\cdot}Tr]_0$ = 17 µM (7.1% of 240 µM)

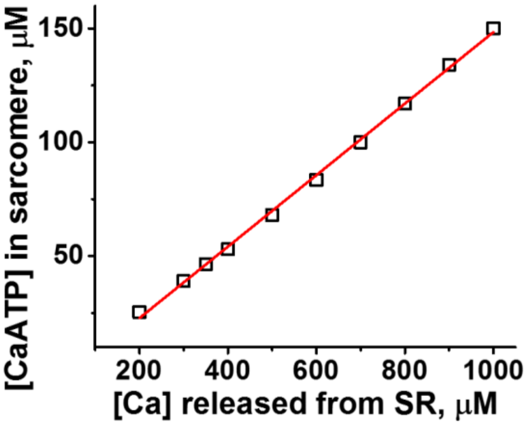


**Figure S 2.** Dependence of maximum [CaATP] transient in sarcomere on calcium, released from the SR. Squares, concentrations calculated in the one compartment model [1]. Line – linear fit to the calculated values. Maximum of [CaATP] in sarcomere linearly depends on the concentration of calcium, released from the SR.

We obtained CaATP transient of the same shape as in Figure 2 of [1]. The dependence of the maximum [CaATP] in the transient on different initial concentrations of calcium, released from the SR is virtually linear (**Figure S *2***). Therefore, we can relate the concentration of CaATP in a sarcomere to the concentration of calcium, released from the SR.

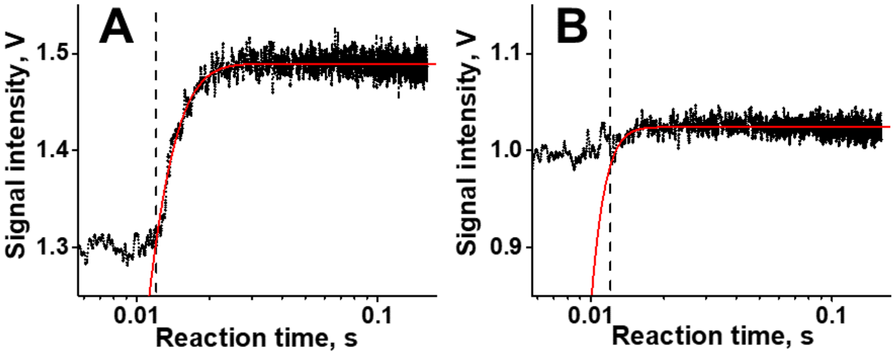


**Figure S 3** MgATP induced actomyosin dissociation, (A) T=12ºC and (B) T=37ºC. Vertical dashed line indicates beginning of the stopped flow kinetics (see **Figure S 4**, calibration of the instrument). In the experiment we observe the flow for 10 ms, then the flow stops. The dead time of the spectrophotometer is 2.6 ms, therefore in this experiment flow stops at 10 ms and we start to fit observed transient from 12.6 ms. Red line is the best fit to a single exponent. At T=37ºC the kinetics of actomyosin dissociation is clearly too fast for reliable determination, the majority of kinetics is hidden in the dead time.

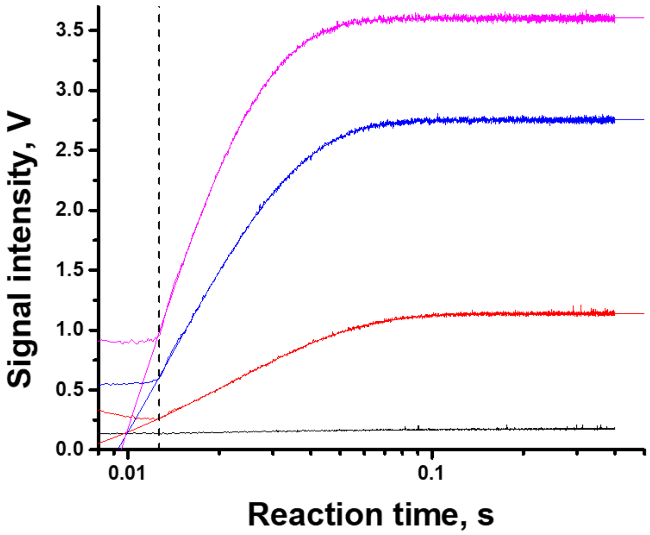


**Figure S 4** Determination of the dead time of the transient spectrophotometer. 100 uM of 8-hydroxyquinoline mixed with 5mM, 2.5mM, 1mM, and 0mM MgCl2 [2]. All concentrations are final. Transients are fitted with single exponential function. Intercept of fitted lines corresponds to the time of the flow stop. Vertical dashed line shows the dead time of the spectrophotometer, 2.6 ms. Flow rate is 8 mL/s per syringe. This calibration gives us information on the parameter $t_0$ in the fitting equation $S(t) = S_o+A\cdot\exp(-k_{obs}\cdot(t-t_0))$, and the time point in the reaction kinetics, we should start the fit (12.6 ms in this case).

### References

1. Baylor, S.M. and S. Hollingworth, *Model of sarcomeric Ca2+ movements, including ATP Ca2+ binding and diffusion, during activation of frog skeletal muscle.* Journal of General Physiology, 1998. **112**(3): p. 297-316.
2. Brissette, P., D.P. Ballou, and V. Massey, *Determination of the dead time of a stopped-flow fluorometer.* Analytical biochemistry, 1989. **181**(2): p. 234-8.
